# RTF: An R package for modelling time course data

**DOI:** 10.1101/2024.06.21.599527

**Authors:** Eva Brombacher, Clemens Kreutz

## Abstract

**Summary:** The retarded transient function (RTF) approach has been introduced as a complementary approach to employing ordinary differential equations (ODEs) for modelling dynamics typically observed for cellular signalling processes. Here, we introduce the R package to the RTF approach, which has originally been implemented within the MATLAB-based *Data2Dynamics* modelling framework. The package enables modelling of time and dose dependencies and includes model reduction to minimize overfitting. It can be applied not only to experimental data but also to trajectories of ODE models, to characterize the dynamics or to identify major targets of experimental perturbations in the low-dimensional representations calculated by the package.

**Availability and Implementation:** The R package RTF is available at https://github.com/kreutz-lab/RTF.

## 1. Introduction

To date, ordinary differential equation (ODE) models are the method of choice in systems biology for mathematically modelling the dynamics of signalling pathway processes within cells. However, ODEs come with some downsides, such as failing to provide analytical solutions and requiring large models and comprehensive amounts of experimental data for the calibration of the unknown model parameters.

The retarded transient function (RTF) modelling approach has been previously introduced as a valuable complement to the traditional ODE-based modelling (Kreutz, 2020). It is tailored to typical dynamic response curves observed in cellular signalling pathways, which often display the following features: new steady states after stimulation, monotone or at most one peak dynamics, and delayed responses. However, it can also be used for time course data not related to signalling, as long as the data displays these features.

The RTF is beneficial in that it allows for fast and simple evaluations and accurate estimates of time- and dose-dependent system dynamics and for a straightforward interpretation of its parameters (amplitudes, rate constants, time constants). It describes the time-dependency of a signalling response by means of a sustained and a transient part, containing, in total, three exponential functions and a nonlinear time transformation, and has previously been extended to also account for dose-dependencies, i.e., the effect of changing treatment doses (Rachel et al., 2024).

Here, we introduce the R (R Core Team, 2024) package for RTF-based modelling, which so far has only been available in MATLAB in the *Data2Dynamics* modelling framework (Raue et al., 2015). It can be applied to experimental time course data as well as to time course data simulated based on ODE models. To prevent overfitting model reduction by stepwise elimination of parameters can be performed. As an additional feature, the package enables the calculation of a low-dimensional representation of the fitted dynamics by employing a uniform manifold approximation and projection (UMAP) on the RTF parameters. This enables a clustering of qualitative differences of the response of signalling compounds. This way, the effects of experimental perturbations, such as knockouts, can be visualized (Fig. 1c).

**Fig. 1:**
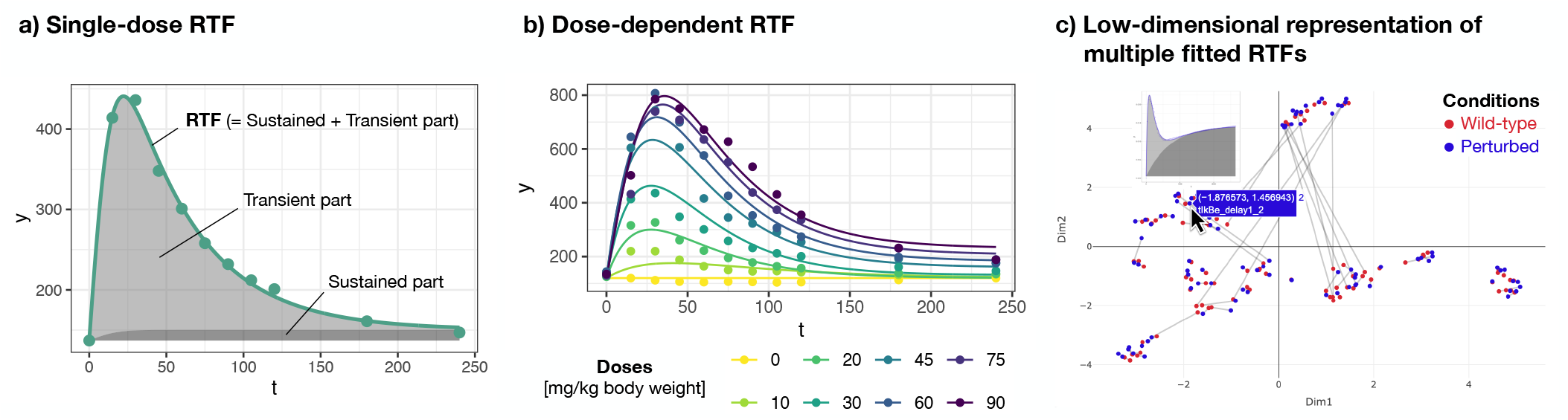
a) Time-resolved data can be modelled by the single-dose retarded transient function (RTF). This subfigure is based on the dataset by Matsumoto et al. (2014) (*matsumoto*), for which plasma amino acid profiles were measured after ingesting different kinds of branched-chain amino acids. The subfigure shows the example of measured leucine plasma levels if 30 mg/kg body weight of leucine was ingested. b) The dose-dependent RTF applied to time-resolved data generated at multiple doses allows. This subfigure is based on the dataset by Matsumoto et al. (2014) (*matsumoto*) and shows the example of measured leucine plasma levels for different leucine concentrations. c) Low-dimensional representation of multiple fitted RTFs. The outcome of multiple fits of time course data can be visualized as a low-dimensional representation calculated from the RTF parameters, where each point represents one time course. This subfigure is based on the ODE model by Almaden et al. (2014), which we used to generate time courses (*almaden*) of the conditions ’Wild-type’ and ’Perturbed’, depicted in red and blue, respectively. To illustrate qualitative changes of the dynamics, the time courses of the same molecular entity for the analyzed conditions are connected by a line. If the time courses of the same molecular entity are too close to each other for the two conditions, the line might not be visible. The cursor indicates the interactive selection of a time course, for which the dynamics should be displayed.

## 2. Retarded Transient Function

The RTF by Kreutz (2020) describes time-dependency as a combination of a sustained and a transient component and is defined as

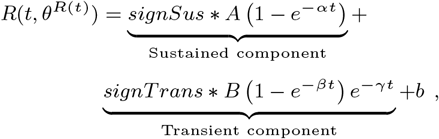

where both components have their individual signs ∈ {−1, 1}*signSus* and *signT rans*, amplitudes *A, B* and the rate constants *α, β*, and *γ* (Fig. 1a). This is complemented by the addition of an offset *b*. The argument *t* is a nonlinear transformation of the experimental time axis with a time-shift parameter *τ*, which accounts for the delayed response of pathway compounds:

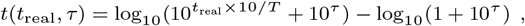

with the real experimental measurement times *t*_real_ and the range of the measurement times *T*. Furthermore, to reduce overfitting, a custom model reduction procedure comprising a stepwise elimination of parameters can be performed.

This approach has recently been extended by describing the dose-dependency of the parameters *A, B, α, β, γ*, and *τ* by Hill equations (Rachel et al., 2024) containing the following three parameters: maximum value *M*, half-maximal quantity *K* (corresponding to EC50), and the Hill coefficient *h*, which accounts for sigmoidality.

The R package presented here incorporates the calculation of the RTF for time- and dose-dependency, hereafter referred to as ‘single-dose RTF’ and ‘dose-dependent RTF’.

## 3. Implementation

The main purpose of the introduced package is to fit the RTF to experimental or simulated time course input data. The input data to be fitted is provided in the form of a data frame containing columns named ’t’ (time) and ’y’ (quantitative value), and (optionally) ‘d’ (dose). In addition, a column ’sigmaExp’ with the standard error of the experimental data can be provided, if it is known. If ’sigmaExp’ is not provided, *σ* will be fitted together with all other parameters.

A central structure for the calculations performed in this package is the *optimObject*, which is a list containing the input data frame with time-resolved data (’data’), and, for the parameters, the vector of the default initial guesses (’initialGuess.vec’) calculated from the data, the vector of the lower bounds (’lb.vec’), the vector of the upper bounds (‘ub.vec’), a vector of the fixed parameters (’fixed’), a boolean vector (’takeLog10’) indicating if log10 is applied to the bounds, the mode in the form of ‘singleDose’ or ‘doseDependent’ (‘modus’), and a vector of the values of the fitted parameters (’fitted’). By applying the RTF() function a list of the stats::optim() (R Core Team, 2024) results for all initial guesses (’optimResults’), the stats::optim() result of the best fit (’bestOptimResult’), and the likelihood value of the best fit (’value’) are added to the optimObject.

For reasons of speed and accuracy of the optimization the analytical derivative of the RTF is calculated, instead of using a finite difference approximation.

The most important implemented functions of the presented R package are:

- *RTF()* estimates the best-fit RTF parameters for the provided input data and can be run in ’singleDose’ (Fig. 1a) or ’doseDependent’ (Fig. 1b) mode, depending on whether signalling data at multiple doses are available. All parameters are jointly estimated based on maximum likelihood by applying multi-start optimization to ensure that the global optimum is found. The sorted multi-start optimization results are visualized in a waterfall plot, where the occurrence of a plateau for the best likelihood value indicates the global optimum. If a user would like to further improve the best fit of a previous result, the previous result can be provided to the function *RTF()* via the argument ’resOld’. The previous result, in turn, becomes complemented with optimization runs from additional initial guesses, with the best fit over the old and new initial guesses being returned.
- *modelReduction()* applies a model reduction (Kreutz (2020)) to the *RTF()* result, i.e., it iteratively eliminates parameters that are according to likelihood ratio tests not required to explain the data. This results in a model with a minimal number of parameters and prevents overfitting.
- *lowDimensionalRTF()* implements the calculation of a low-dimensional representation of multiple fitted RTFs (Fig. 1c). Specifically, the RTF parameters are estimated for multiple time courses and based on those parameters a two-dimensional UMAP is generated. If the dynamics of certain molecular species differ between two conditions, this should be reflected in changes in the RTF parameters, and, in turn, in changes in the position within the UMAP. The formation of a cluster in the UMAP indicates that, based on the RTF parameters, the dynamics are similar for the molecular species of the respective cluster.
- *plotInteractiveUMAP()* generates an interactive UMAP plot based on the estimated RTF parameters for multiple time courses, where for each time course (represented by a point), its dynamics are displayed upon mouseover. The example in Fig. 1c shows the UMAP for the NF*κ*B pathway dynamics of the dataset by Almaden et al. (2014) (introduced below), where two different conditions are compared and the time courses of the same molecular species are connected by a line.
- *plotData()* plots the input data to be fitted. For dose-dependent datasets, the data points are color-coded according to the respective dose.
- *plotRTF()* plots the *RTF()* results as illustrated in Fig. 1a and b.

Example datasets can be simulated using the function *getSimData()* for modes ‘singleDose’ or ‘doseDependent’. In addition, the following datasets are made available in our R package:

- *matsumoto*: Experimental dataset by Matsumoto et al. (2014), where the plasma profiles of 21 amino acids were investigated after ingesting 10–90 mg/kg body weight of branched-chain amino acids (leucine, isoleucine, or valine). Data is available for 11 different time points.
- *strasen*: Simulated dataset based on the cell class model by Strasen et al. (2018b), which reflects six signalling classes observed upon stimulation with 100 pM Transforming growth factor (TGF) *β*1. For six cell classes the time courses of 23 signalling proteins were simulated using JWS Online (Strasen et al., 2018a; Olivier and Snoep, 2004). Data is available for 101 different time points.
- *almaden*: Simulated protein dataset based on the model of the NF-*κ*B-signalling system in B cells introduced by Almaden et al. (2014). This time course dataset is modelled using ‘Data2Dynamics’ (Raue et al., 2015) and contains simulated time course data for 91 molecular entities. This time course data has been generated for two conditions in the form of a genetic perturbation that prevents the processing of p100 to p52 (condition 2) and wild-type controls (condition 1), resulting in in total 182 time courses. Data is available for 101 different time points.

A basic example for applying the RTF R package may look as follows:

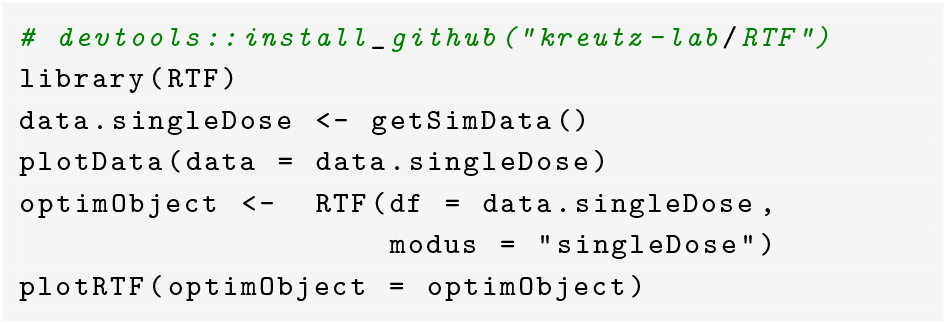

## 4. Discussion

The introduced R package makes the RTF approach – a novel modelling approach for time- and dose-dependent responses typically observed for signalling pathway entities – accessible to the large R community in bioinformatics.

Applying the RTF results in few parameters characterizing the dynamics which offers a further opportunity to interpret signalling dynamics and the effect of perturbations. By calculating a two-dimensional representation, the package indicates compounds with similar dynamics, e.g., by means of clustering. In addition, comparing multiple experimental or biological conditions in such representations can reveal changing dynamic behavior, indicated, for instance, by compounds switching from one cluster to another under a perturbed condition.

In the future, the R package could be extended such that the uncertainty of parameters will be calculated.

## 5. Acknowledgements

The authors sincerely thank Stefan Legewie for his insightful suggestions.

## 6. Conflict of interest

None declared.

## 7. Funding

This work was supported by the Deutsche Forschungsgemeinschaft (DFG, German Research Foundation) under Germany’s Excellence Strategy (CIBSS-EXC-2189-2100249960-390939984) (C.K. and E.B.).

## Notes

### Competing Interest Statement

The authors have declared no competing interest.

## References

J. V. Almaden, R. Tsui, Y. C. Liu, H. Birnbaum, M. N. Shokhirev, K. A. Ngo, J. C. Davis-Turak, D. Otero, S. Basak, R. C. Rickert, et al. A pathway switch directs BAFF signaling to distinct NFκB transcription factors in maturing and proliferating B cells. Cell reports, 9(6):2098–2111, 2014.

C. Kreutz. A new approximation approach for transient differential equation models. Frontiers in Physics, 8:1–14, 2020.

T. Matsumoto, K. Nakamura, H. Matsumoto, R. Sakai, T. Kuwahara, Y. Kadota, Y. Kitaura, J. Sato, and Y. Shimomura. Bolus ingestion of individual branched-chain amino acids alters plasma amino acid profiles in young healthy men. Springerplus, 3:1–13, 2014.

B. G. Olivier and J. L. Snoep. Web-based kinetic modelling using JWS Online. Bioinformatics, 20(13):2143–2144, 2004.

R Core Team. R: A Language and Environment for Statistical Computing. R Foundation for Statistical Computing, Vienna, Austria, 2024. URL https://www.R-project.org/.

T. J. Rachel, E. Brombacher, S. Woehrle, O. Gross, and C. Kreutz. Dynamic modelling of signalling pathways when odes are not feasible. bioRxiv, 2024. doi: 10.1101/2024.04.18.590024. URL https://www.biorxiv.org/content/early/2024/04/20/2024.04.18.590024.

A. Raue, B. Steiert, M. Schelker, C. Kreutz, T. Maiwald, H. Hass, J. Vanlier, C. Tönsing, L. Adlung, R. Engesser, et al. Data2Dynamics: a modeling environment tailored to parameter estimation in dynamical systems. Bioinformatics, 31(21):3558–3560, 2015.

J. Strasen, U. Sarma, M. Jentsch, S. Bohn, C. Sheng, D. Horbelt, P. Knaus, S. Legewie, and A. Loewer. JWS Online: strasen2018 Fig5A. https://jjj.bio.vu.nl/models/experiments/strasen2018_fig5a/, 2018a. Accessed: 2023-11-19.

J. Strasen, U. Sarma, M. Jentsch, S. Bohn, C. Sheng, D. Horbelt, P. Knaus, S. Legewie, and A. Loewer. Cell-specific responses to the cytokine TGF β are determined by variability in protein levels. Molecular systems biology, 14(1):e7733, 2018b.

